# Computational analysis of molecular networks using spectral graph theory, complexity measures and information theory

**DOI:** 10.1101/536318

**Authors:** Chien-Hung Huang, Jeffrey J. P. Tsai, Nilubon Kurubanjerdjit, Ka-Lok Ng

## Abstract

Biological processes are based on molecular networks, which exhibit biological functions through interactions among the various genetic elements. This study presents a graph-based method to characterize molecular networks by decomposing them into directed multigraphs: network motifs. Spectral graph theory, reciprocity, and complexity measures were utilized to quantify the network motifs. It was found that graph energy, reciprocity, and cyclomatic complexity can optimally specify network motifs with some degree of degeneracy. A total of 72 molecular networks were analyzed, of three types: cancer networks, signal transduction networks, and cellular processes. It was found that molecular networks are built from a finite number of motif patterns; hence, a graph energy cutoff exists. In addition, it was found that certain motif patterns are absent from the three types of networks; hence, the Shannon entropy of the motif frequency distribution is not maximal. Furthermore, frequently found motifs are irreducible graphs. These are novel findings: they warrant further investigation and may lead to important applications.

The present study provides a systematic approach for dissecting biological networks. Our discovery supports the view that there are organizational principles underlying molecular networks.

## Background

### Biological networks, network motifs, and graphlets

Molecular networks are the basis of biological processes, in which biological functions emerge through interactions among the various genetic components. A network can be modeled by a collection of smaller modules; each module is expected to perform specific functions, and is separable from the functions of other modules [1-3]. Such modular networks can be decomposed into smaller units, known as network motifs. These motifs show interesting dynamical behaviors, in which cooperativity effects between the motif components play a critical role in human diseases.

We classify network-based analysis into the following major categories: (1) motif identification and analysis, (2) global architecture study, (3) local topological properties, and (4) robustness of the network under different types of perturbations. For the first category, there are a number of publicly available network motif detection tools namely, MFINDER [4], MAVISTO [5], FANMOD [6], NetMatch [7], and SNAVI [8].

For the second category, many studies have employed random graph theory to characterize the global structure of molecular networks: for example, whether a network is assortative or has the small-world property [9-10]. For instance, it has been shown that protein-protein interaction networks are scale-free or described by hierarchical network model [11].

For the third category, topological graph theory has been utilized to characterize networks by computing topological parameters, such as betweenness centrality, closeness centrality, clustering coefficients, and eigenvector centrality [12-16].

For the last category, it has been shown that molecular networks are robust under random perturbation but fragile under attack perturbation [17]. Further work has demonstrated that molecular networks are also fragile under degree-based, betweenness-based, and brokering coefficient-based perturbations [18].

Besides network motif description, Przulj [19-20] utilized a graphlet-based approach to examine the network comparison problem.

It was demonstrated that directed graphlets are superior for comparing directed networks [21] and they are effective for studying brain networks [22].

Our study focuses on networks composed of regulatory interactions, such as gene regulation networks and signal transduction networks but not protein-protein interaction networks (undirected graphs). We work with network motifs directly; therefore, our method differs from the graphlet approach.

Although many published works exist on network analysis, many important issues still remain to be investigated. Most previous studies have utilized graph metrics to analyze network topology, and so a very relevant question remains unanswered: do these topological parameters convey enough knowledge about the networks? The answer seems to be negative. Little is known about the architectures or organizational principles of molecular networks. For instance, can we have a unique label for different motifs? Do certain motif patterns occur in a network at a higher frequency?

Seminal works on the use of the concepts of information content, topology, and entropy in biology were carried out by Dancoff & Quastler [23], Rashvesky [24-25], and Mowshowitz [26-27]. In particular, Mowshowitz presented an entropy-based method to measure the complexity of a graph by decomposing it into equivalence classes.

In this study, it is hypothesized that network motifs are the fundamental building blocks of a network. In other words, motifs are treated as the core components of a network. This is similar in spirit to the work of Mowshowitz [27]. Therefore, we propose that network properties are captured by motifs comprising *N* nodes, which are referred to as *N*-node motifs in the following discussion. To systematically characterize a large network, one identifies the 3-node motifs, 4-node motifs, up to the *N*-node motifs embedded in the network.

For a directed graph, a total of 2, 13, 199, 9364, and 1530843 possible patterns can be defined for the 2-node, 3-node, 4-node, 5-node, and 6-node motifs, respectively [28-29]. Since the problem of identifying *N*-node motifs in a large network is NP-complete [30], we worked with 3-node motifs and 4-node motifs only. Motifs composed of five or more nodes are neglected as a first approximation. As we explain below, this approximation could provide useful insights into dissecting the design principles underlying molecular networks. Motifs composed of five or more nodes will be considered in future study.

An earlier work [31] has shown that certain motifs do not appear significantly more frequently than those appearing in corresponding random graphs; nevertheless, those motifs still play functional roles. This justifies our approach because the present work identifies all possible 3-node and 4-node motifs, regardless of their frequency of occurrence. In other words, we adopt the notion that motifs are the basic building blocks but do not necessarily occur frequently in a network.

Adami [32] studied undirected colored graphs (in which nodes are labeled with different colors) and showed that the relative frequency of the colored motifs can be used to define the information content of the network. In the present work, we consider motifs that are *directed* graphs and could possibly contain cycles.

### Spectral graph theory, reciprocity, complexity measures, and information theory

To characterize network motifs, we utilized the following concepts: spectral graph theory (SGT), reciprocity, and complexity measures. SGT is a powerful approach that has been applied in many areas, including computer science and computational biology [33-34]. The eigenvalues of a matrix defined on a graph play an essential role in inferring the structural properties of the graph [35]. According to Mowshowitz [36], the characteristic polynomial of the adjacency matrix of a graph distinguishes between non-isomorphic graphs. Reciprocity is a parameter that quantifies the degree of bidirectional connection of a network motif.

Complexity arises from the interactions among the constituent components. Many complexity measures have been proposed, but there is no standard or formal definition of complexity metrics that can be applied in all circumstances. Each complexity measure has strengths and weaknesses [37]. Early work on defining complexity for directed graphs and infinite graphs can be traced back to Mowshowitz [38]. The concept of graph complexity indices has been applied to infer the hierarchical order of chemical structures [39]. Given a network motif pattern, we make use of two commonly used complexity measures to characterize the motif.

It is possible that some of the network motifs are associated with the same graph energy (degenerated motifs). Wilson & Zhu [40] have proposed to combine the spectra of two graph matrices to reduce the cospectrality problem for undirected graphs and trees. Their results showed that their method can reduce the number of cospectral pairs of graphs but they are still not completely distinguishable. In addition, graph descriptors are a useful concept to classify complex networks [41]. In this study, we used a greedy algorithm to search for an optimal set of parameters that maximize the removal of degenerate motifs. The parameters we suggested include not only the motif spectrum but also the graph energy, reciprocity, and complexity measures.

The concept of information entropy has been applied extensively in cancer biology studies. For instance, it was reported that cancer networks exhibit high information entropy [42], as well as increased network entropy [43] and signaling entropy [44]. We make use of entropy to measure the frequency distributions of the occurrence of motifs for the three types of molecular network.

In our previous work [45], we already laid a foundation for the present study. In another recent work [46], we have extended the previous work [45] by developing a motif finding algorithm, *PatternFinder,* to identify the 3-node motifs and 4-node motifs in cancer networks, signal transduction networks, and cellular processes.

## Methods

### Workflow of present study

Figure 1 depicts the workflow of the present study.

**Figure 1.**
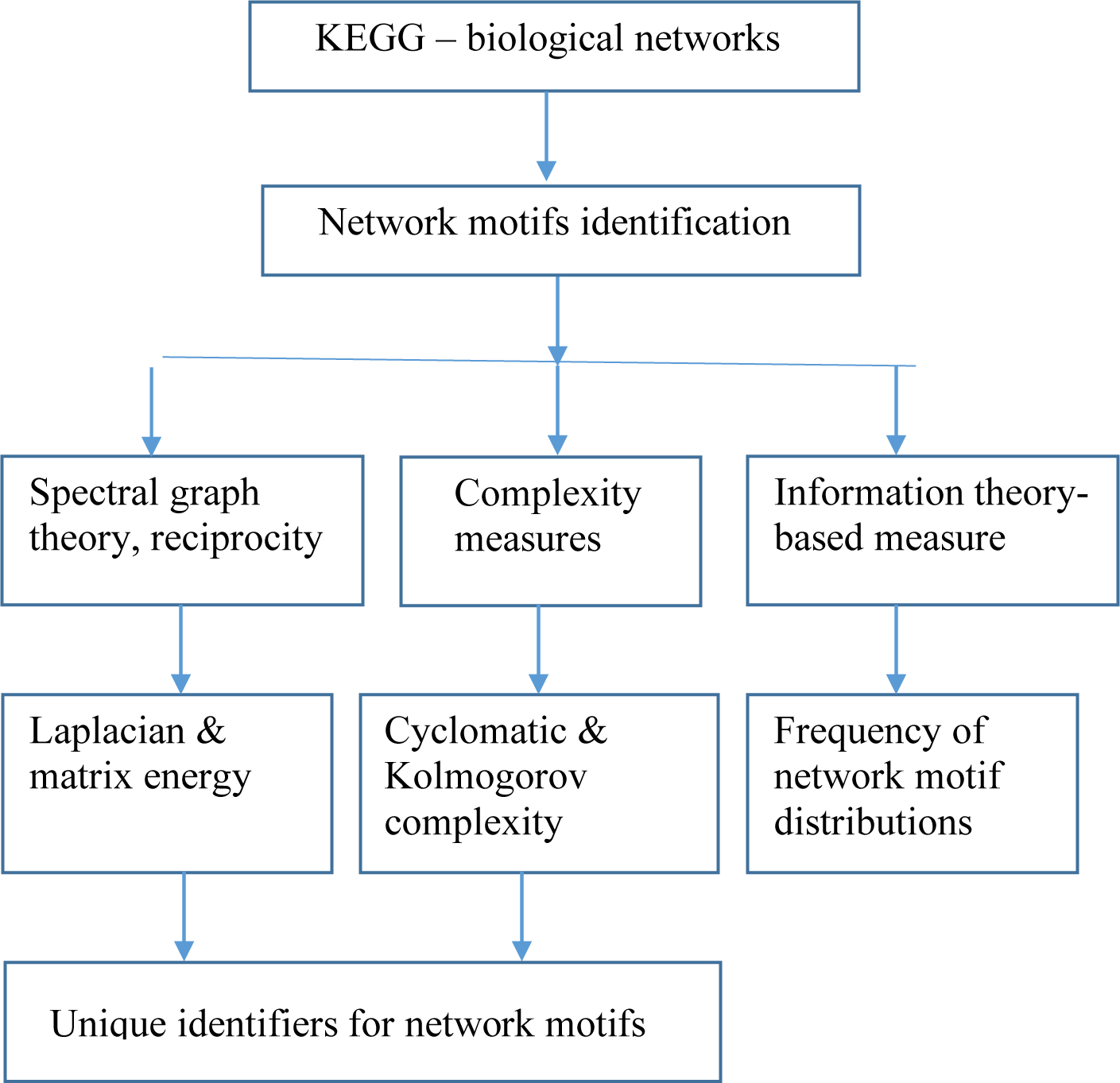
The workflow of the present study.

### Input data

Network information was obtained from the KEGG database (August 2017) [47]. Four families of networks were employed, including: (i) Environmental Information Processing, (ii) Cellular Processes, (iii) Organismal Systems, and (iv) Human Cancers.

Not every network recorded by KEGG was imported. After inspection, we disregarded networks composed of several separate components, such as the “Two-component system,” “MicroRNAs in cancer,” “Chemical carcinogenesis,” and “Viral carcinogenesis”. In addition, we combined the networks labeled with the name “signaling pathway,” and called them “signal transduction networks (STNs)”. We note that STNs range across different families in the KEGG classification, including “Signal transduction,” “Immune system,” and “Endocrine system”.

In total, we collected 17 cancer networks, 46 STNs, and 9 cellular processes. We downloaded KEGG pathway KGML files and made use of the KEGGScape [48] and KEGGparser [49] packages to visualize and save the node and edge information for each network.

In Supplementary File 1, Supplementary Table S1 summarizes the nodes, edges, and motif-associated node information for the 17 cancer networks. The complete list of node and edge information of the 46 STNs and 9 cellular processes can be found in Supplementary Table S2 and Supplementary Table S3, respectively, in the same file.

### Adjacency matrix

By analyzing the connectivity of each gene, one constructs an adjacency matrix, ***A***, to represent the interaction network. In total, there are 13 3-node motifs and 199 4-node motifs [3,50].

It is possible that some motifs are subgraphs of other motifs (structural motifs). In a previous work [51], such subgraphs are called functional motifs. In a brain network, a structural motif and functional motif represent an anatomical building block and the elementary processing mode of a network, respectively.

We have developed an algorithm named *PatternFinder* to enumerate all possible functional motifs embedded in the 3-node motifs and 4-node motifs. Details of *PatternFinder* are given in Supplementary File 1 – Supplementary Table S4.

### Characterization of network motifs:graph energy, reciprocity and graph complexity

The energy of a graph is an invariant [52-54], and is equal to the sum of the absolute values of the eigenvalues of the adjacency matrix ***A***. Originally, the concept of graph energy introduced by Gutman was applied to study undirected graphs and has been applied to estimate the π–electron energy of hydrocarbons.

The adjacency matrix ***A*** can be expressed in terms of its eigenvectors and eigenvalues. Since ***A*** is a nonsymmetric matrix in general, its eigenvalues may be complex and all of its eigenvectors are nonorthogonal. Let *n, e*, and *d*_*i*_ denote the number of nodes, number of edges, and degree of the *i*th node of graph *G*, respectively; *G* is called an (*n, e*)-graph. The energy of a graph *G, E(G)*, is defined by

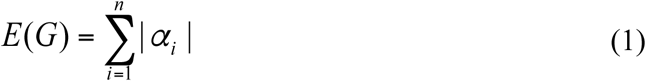

where *α*_*i*_ denotes the *i*th eigenvalue of ***A***. The sum of all of the eigenvalues is always equal to zero.

Assume that the graph energy eigenvalues are labeled in descending order: that is, α_1_ ≥ α_2_ ≥ … ≥ α_*n*_, while the whole spectrum is denoted by *Sp*(*G*) = [α_1_, α_2_, … α_n_]. The largest eigenvalue is referred to as the spectral radius of graph *G* [55].

In spectral graph theory, there are two other matrices—Laplacian [56] and signless Laplacian [57-58]—that can be defined to characterize graphs. The Laplacian matrix ***L*** and signless Laplacian matrix ***Q*** of a graph *G* are defined as ***L*** = ***D*** – ***A*** and ***Q*** = ***D*** + ***A*** respectively, where ***D*** is a diagonal matrix in which the diagonal elements are the node degrees. The Laplacian energy of a graph *G, LE*(*G*), is defined by

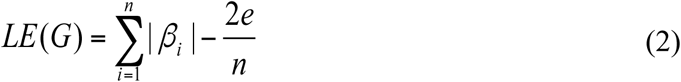

where | *β*_*i*_ | denotes the absolute value of the *i*th eigenvalue of ***L***. There is an analogy between the properties of *E*(*G*) and *LE*(*G*), but some significant differences remain between these two quantities [59].

The signless Laplacian energy of graph *G, QE*(*G*), is defined by

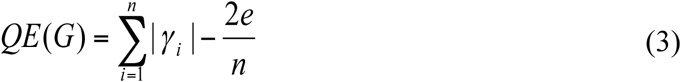

where |γ_*i*_| denotes the absolute value of the *i*th eigenvalue of ***Q***.

A more general definition of graph energy was suggested by Nikiforov [60-61]. Let ***M*** be an *n* × *n* real matrix and the singular values be denoted by *s*_1_, *s*_*2*_, … *s*_*n*_. The singular values of ***M*** are equal to the positive square roots of the eigenvalues of ***MM***^***t***^, where ***t*** denotes matrix transpose. Let ***M*** equal ***A, L***, or ***Q*** and consider the eigenvalues of ***AA***^***t***^, ***LL***^***t***^, and ***QQ***^***t***^. The total energy, ***ME***, obtained from ***M***, is defined by

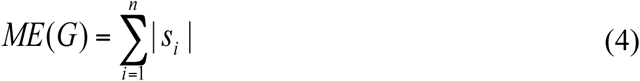

*ME*(*G*) is called generalized energy. We extend the definition to consider matrix products of the form ***MN***^t^, and therefore define three additional energies: ***AL***^***t***^, ***AQ***^***t***^, and ***LQ***^t^. We call these *asymmetric* generalized energies. The sums of the absolute values of the eigenvalues of ***MM***^t^and ***M***^t^***M*** are the same. This also holds for ***MN***^t^ and ***NM***^t^. Therefore, one needs to compute ***MM***^t^ and ***NM***^t^ only. The advantages of using *asymmetric* generalized energies will be demonstrated later in this article. To the best of our knowledge, no (or few) previous studies have made use of *asymmetric* generalized energies to characterize network motifs. In total, we have devised nine graph energies to describe the motifs. We also note that Adiga et al. [62] proposed a parameter named skew energy, obtained from the skew-adjacency matrix, to characterize directed graphs; however, this parameter does not apply to graphs consisting of multiple arcs (multigraphs).

Several studies [63-64] have suggested that reciprocal links in directed networks play an important role in dynamical processes and network growth. The traditional definition of reciprocity is *R = L*^↔^ */ L*, where *L*^↔^ and *L* denote the number of “edges pointing in both directions” and the total number of edges respectively. This definition of reciprocity was modified by Garlaschelli and Loffredo [63], who defined reciprocity *r* as the correlation coefficient between the entries of the adjacency matrix, ***A***, given by

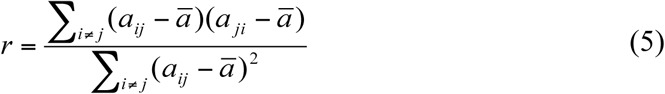

where *a*_*ij*_ equals one if there is an edge from node *i* to node *j*; the average, ā, is defined by

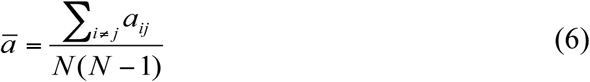

A positive value of *r* indicates that the motif has bidirectional connections, whereas a negative *r* implies that the motif has either an in-connection or out-connection.

To further understand the connectivity structure of network motifs, we seek metrics that can be used to measure graph complexity. In software engineering, the cyclomatic complexity (*CC*) is a metric developed by McCabe [65] to measure the complexity of a program by using its control flow graph. *CC* is defined by the expression *CC* = *e – N + 2P*, where *e* and *N* denote the number of edges and number of nodes of the graph, and *P* denotes the number of predicate/exit nodes [37,65]. Node and edge denote a program unit and the execution order of the program. *CC* depends only on the global decision structure (the number of edges and nodes) of a program.

In addition to *CC*, we utilize the algorithmic complexity measure, the Kolmogorov complexity (*KC*), to characterize graph complexity. Essentially, the *KC* of a bit string is given by the smallest computer program that can generate the string. Given the adjacency matrix (or the equivalent bit string), we use the block decomposition method (BDM) [66] to determine the *KC* for both 3-node [67] and 4-node motifs. A bit string with a high *KC* has a higher degree of randomness, contains more information, and is less compressible. A complete graph has a smaller *KC* value, whereas a random graph has higher *KC* and is less compressible.

### Unique identifiers for network motifs

Every 3-node motif and 4-node motif has a different *KC* value, so the *KC* can be used as a unique identifier. However, given the graph energy, asymmetric graph energies, graph energy spectrum, reciprocity, and *CC*, we seek to determine a minimal set of parameters that can serve as a label of the network motifs. This set of parameters describes certain aspects of the motifs differently than the algorithmic complexity measure. To the best of our knowledge, the concept of using energy, reciprocity, and *CC* in labeling network motifs is novel. The pseudocode for determining the minimal set of parameters is based on a greedy strategy and is described in Supplementary File 1 – Supplementary Table S5.

### Frequently found motifs, network entropy, and network similarity

Given a molecular network, *PatternFinder* identifies both the sets of 3-node motifs and 4-node motifs. Two motifs with the same ID may partially embed the same genetic element(s); these two motifs are counted twice in our calculations. We expect that certain motif patterns that occur with higher probabilities are the dominant underlying network structure. Let 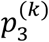 denote the frequency (probability) distribution of a 3-node network motif, where *k* denotes one of the 13 patterns. The Shannon entropy for 3-node motifs and 4-node motifs, *H*_*3*_ and *H*_*4*_, of a molecular network are computed. The normalized Shannon entropies for the 3-node motifs and 4-node motifs are given by *H*_*3R*_ = *H*_*3*_ / log_2_(13) and *H*_*4R*_ = *H*_*4*_ / log_2_(199), respectively.

## Results

Given the 3-node motifs and 4-node motifs, we used *PatternFinder* to identify their subgraphs (all possible functional motifs). For the 3-node motifs, it was found that motif “id_6” (SIM), motif “id_12” (cascade), and motif “id_36” (MIM) are not composed of any 3-node functional motifs. For the 4-node motifs, there are eight motifs that are not composed of any 4-node functional motifs: motif “id_14” (SIM), motif “id_28,” motif “id_74,” motif “id_76” (MIM), motif “id_280,” motif “id_328” (cascade), motif “id_392,” and motif “id_2184”. These eight motifs exhibit the property of *irreducibility.* However, each one of the eight motifs is embedded with exactly one 3-node functional motif. In other words, given the 4-node motifs, the *irreducible* property does not apply if we consider motifs composed of three nodes. Supplementary File 2 summarizes the functional motifs for 3-node motifs, 4-node motifs, and 3-node motifs embedded in 4-node motifs, where integers “1” and “0” denote the presence or absence of a functional motif, respectively.

### Spectral graph theory, reciprocity, and complexity measures

Table 1 summarizes the results of the nine graph energies and edge information for the 3-node motifs. First, since some of the matrices, such as *L* and *Q*, are asymmetric, their eigenvalues are complex in general. In fact, among the 3-node motifs, motif “id_98” has a pair of complex conjugate eigenvalues, and their associated eigenvectors are composed of complex components.

**Table 1.**
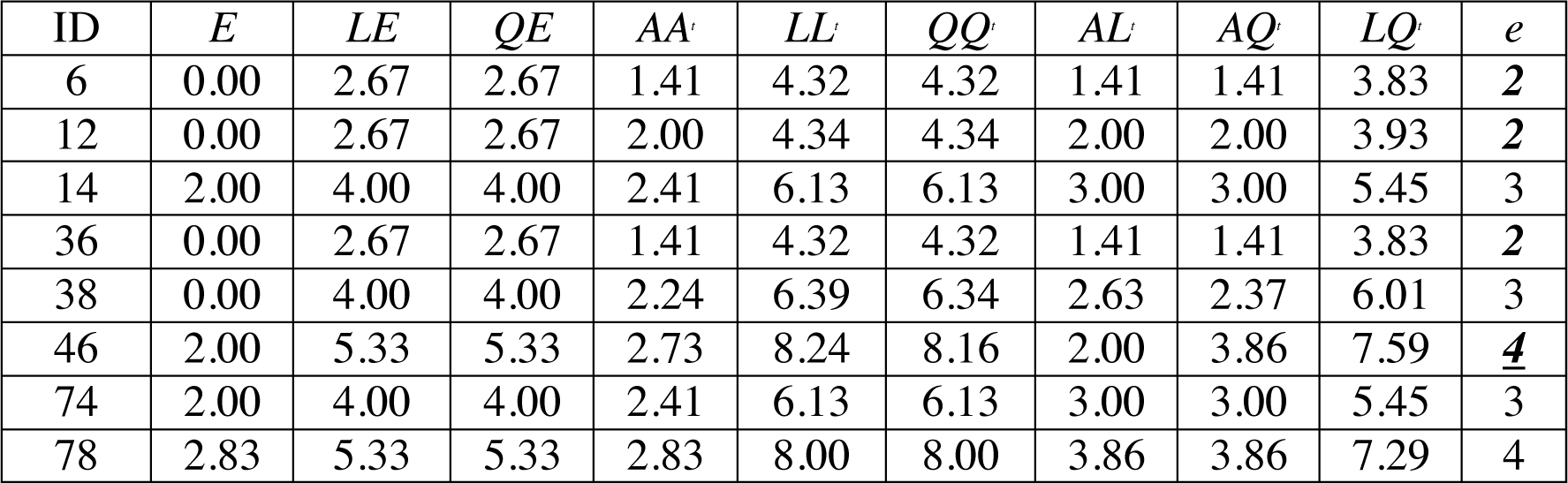
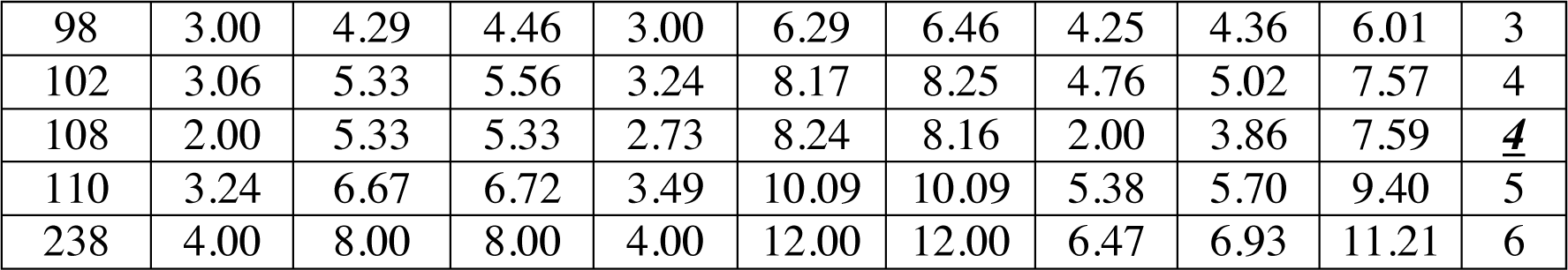
The results of the nine graph energies and edge information for the 3-node motifs.

Second, graph energy is correlated with the number of edges of a motif. For instance, the graph energies of fully connected 3-node motifs and 4-node motifs are maximal, despite having different energy definitions.

Third, it is quite common for certain motifs to have the same graph energy; that is energy-degenerated motifs are rather common. Two motifs are said to be *equienergetic* if they have the same total energy. For instance, two pairs of motifs (“id_6” and “id_36,” and “id_14” and “id_74”) are *equienergetic*, regardless of the graph energy definition. The results of the 4-node motif graph energies and eigenvectors are given in Supplementary File 3.

Fourth, although the results of the nine graph energies are quite similar, there are differences among them: for instance, the multiplicity of the energy levels is somewhat different. For the 3-node motifs, the multiplicities of graph energy *E,* 0, 2, and 2.83 are 4, 4, and 1, respectively. For *QE*, there are three energy values, 2.67, 4.00, and 5.33, that are associated with the multiplicity of 3, 3, and 3, respectively.

Fifth, energy-degenerated motifs may or may not have identical spectra, *Sp*(*G*). This suggests that the use of *Sp*(*G*) could allow for further distinction between the motifs. More details are given below in the “Unique identifiers for network motifs” section.

In Supplementary File 1, Supplementary Table S6 summarizes the lower (*E*_*min*_) and upper (*E*_*max*_) graph energy bounds and ratios for the 3-node motifs and 4-node motifs. For the 3-node motifs, the ratios are bounded between 2 and 4.91. These ratios are slightly larger for 4-node motifs: they are bounded between 3.00 and 6.88. We found that most of the molecular biological networks are not composed of motifs with large graph energies; therefore, the maximum ratio cannot be achieved. Details are reported below in the “Network motifs absent from the network” section.

### Reciprocity of motifs

Table 2 depicts the traditional reciprocity *R*, reciprocity *r*, and ā for the 3-node motifs. Most of the *R* values are zero, which indicates that there is no edge pointing in both directions. Positive and negative values of *r* denote the presence of cycles. Of the 13 reciprocity values, nine are negative, meaning that the majority of the 3-node motifs have either in-connections or out-connections only. We note that motifs containing one or two cycles can still have negative reciprocity values. The complete sets of *R, r*, edges and ā values of the 4-node motifs are given in Supplementary File 4.

**Table 2.** The results of traditional reciprocity (*R*), reciprocity (*r*), edge (*e*) and average reciprocity (ā) of the 3-node motifs.

| ID | $R$ | $r$ | $e$ | $\bar{a}$ |
| --- | --- | --- | --- | --- |
| 6 | 0 | -0.5 | 2 | 1/3 |
| 12 | 0 | -0.5 | 2 | 1/3 |
| 14 | 0 | 1/3 | 3 | 0.5 |
| 36 | 0 | -0.5 | 2 | 1/3 |
| 38 | 0 | -1 | 3 | 0.5 |
| 46 | 0 | -0.5 | 4 | 2/3 |
| 74 | 0 | 1/3 | 3 | 0.5 |
| 78 | 1 | 1 | 4 | 2/3 |
| 98 | 0 | -1 | 3 | 0.5 |
| 102 | 0 | -0.5 | 4 | 2/3 |
| 108 | 0 | -0.5 | 4 | 2/3 |
| 110 | 0 | -0.2 | 5 | 5/6 |
| 238 | 1 | 1 | 6 | 1 |

### Graph complexity: cyclomatic complexity and Kolmogorov complexity

For the 3-node motifs, Table 3 summarizes the results of the cyclomatic complexity (*CC*) and Kolmogorov complexity (*KC*), and their rankings. The ranges of these *CC* and *KC* values are 0–3 and 23.34–25.50, respectively. The complete sets of *CC* and *KC* values of the 4-node motifs are given in Supplementary File 5, where the ranges of *CC* and *KC* values are 0–8 and 33.80–43.74, respectively. These findings are compatible with the notion that motifs composed of more nodes have higher complexity.

**Table 3.** The results of the cyclomatic complexity (*CC*) and Kolmogorov complexity (*KC*), and their ranking, for the 3-node motifs.

| ID | *CC* | *KC* | *Rank of CC* | *Rank of KC* |
| --- | --- | --- | --- | --- |
| 6 | 3 | 23.34 | 11 | 1 |
| 12 | 1 | 23.83 | 3 | 3 |
| 14 | 2 | 24.30 | 8 | 6 |
| 36 | 1 | 23.55 | 3 | 2 |
| 38 | 2 | 24.87 | 8 | 8 |
| 46 | 3 | 25.50 | 11 | 13 |
| 74 | 0 | 23.85 | 1 | 4 |
| 78 | 1 | 25.00 | 3 | 9 |
| 98 | 0 | 24.82 | 1 | 7 |
| 102 | 1 | 25.01 | 3 | 10 |
| 108 | 1 | 25.11 | 3 | 11 |
| 110 | 2 | 25.25 | 8 | 12 |
| 238 | 3 | 24.14 | 11 | 5 |

A network motif with a large *CC* value suggests a more complex decision structure. From Table 3, it is apparent that *KC* can serve as a parameter for distinguishing motif patterns without any degeneracy. In other words, no two motifs have the same *KC*; this is also true for the 4-node motifs. Motif “id_238” is a complete graph that is described by the binary string “011101110,” and this string corresponds to lower algorithmic complexity (5^th^ rank).

Next, we examined the correlations between the two complexity measures. We ranked *CC* and *KC* in ascending order and computed their Spearman Rank Correlation Coefficients (*SRCC*). The correlation is not perfect; for example, motif “id_238” is associated with the largest *CC* value (rank), but this is not the case for *KC* (5^th^ rank). *CC* and *KC* show a relatively weak correlation—0.083 and 0.381—at the 3-node and 4-node levels, respectively. This is because *CC* and *KC* have different meanings: *CC* measures the complexity of a motif’s decision structure (the number of independent gene regulation paths), while *KC* is an algorithmic measure which characterizes the randomness and compressibility of a bit string.

Lastly, we investigated whether graph energy is proportional to graph complexity. The results are listed in Supplementary File 1 – Supplementary Table S7. *KC* exhibits a modest correlation with all the graph energies at the 3-node and 4-node levels. In contrast, *CC* exhibits relatively weak and modest correlations with graph energy, at both the 3-node and 4-node motif levels.

Supplementary File 1 – Supplementary Table S8 summarizes the results of strength of *SRCC* (including minimum, maximum, and ranges) between graph complexity and graph energy for 3-node and 4-node motifs. Our results suggest that there are relatively weak (3-node *CC*) and modest correlations (3-node motif *KC* and 4-node motif *CC and KC*) between graph complexity and graph energy.

### Unique identifiers for network motifs

This section reports the results of determining an optimal parameter combination that maximizes the removal of degenerated motifs. As shown in Table 4, three cases are considered. “Case a” makes use of graph energy only, “case b” utilizes graph energy *r* and *CC*, and “case c” employs energy, *r, CC*, and the energy spectrum. After including *r* and *CC*, we can distinguish more motifs. The use of *AL*^*t*^, *r, CC*, and energy spectrum can fully distinguish the 3-node motifs. For 4-node motifs, the use of *LL*^*t*^, *QQ*^*t*^ and *LQ*^*t*^ achieves the best level of distinguishability: 136 out of 199 (68.3%). Compared with *E, LE*, and *QE*, both *symmetric* and *asymmetric* generalized energies serve as superior measures for distinguishing different motif patterns.

**Table 4.**
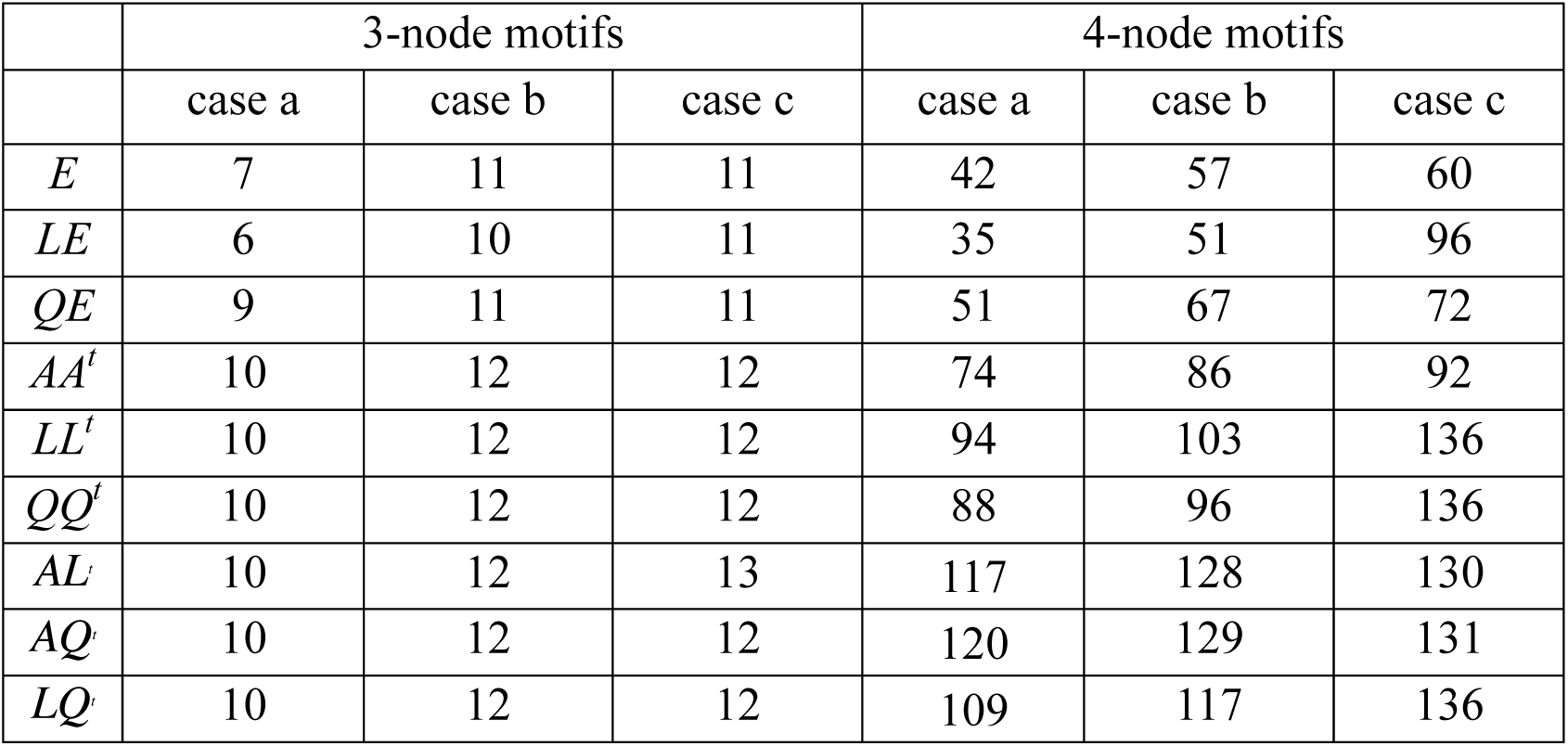
The number of distinguishable motifs using optimal parameter combination of graph energy, *r, CC*, and energy spectrum. “Case a” uses graph energy only, “case b” uses graph energy, *r*, and *CC*, and “case c” uses energy, *r, CC*, and graph energy spectrum.

### Frequently found network motifs

Among the 17 cancer networks, 46 STNs, and nine cellular processes, there are 15 (88.2%), 40 (87.0%), and 7 (77.8%) networks, respectively, where more than 70% of nodes are embedded in both 3-node motifs and 4-node motifs. Therefore, motif-associated nodes account for a major portion of each network.

To determine frequently occurring motifs, we tabulate the frequency of occurrence of each motif pattern, normalize the frequency to one, and compute the average normalized frequency (probability) across the studied networks. Table 5 summarizes the top seven most frequently found 3-node motifs and 4-node motifs in cancer networks.

**Table 5.** The top seven most frequently found 3-node motifs and 4-node motifs in cancer networks. SIM denotes simple input module, MIM denotes multiple input module, and FFL denotes feed-forward loop.

| | ID | average<br>probability | reciprocity<br>$r$ | Name, embedded motif ID |
| --- | --- | --- | --- | --- |
| 3-node |  |  |  |  |
| 1 | 6 | 0.421 | -1/2 | SIM |
| 2 | 12 | 0.414 | -1/2 | Cascade |
| 3 | 36 | 0.152 | -1/2 | MIM |
| 4 | 38 | 0.0091 | -1 | FFL, 12, 36 |
| 5 | 74 | 0.0022 | 1/3 | 12 |
| 6 | 14 | 0.0016 | 1/3 | 6 |
| 7 | 98 | 0.00092 | -1 | 3-cycle, 12 |
| 4-node |  |  |  |  |
| 1 | 14 | 0.224 | -1/3 | SIM |
| 2 | 328 | 0.158 | -1/3 | Cascade |
| 3 | 28 | 0.148 | -1/3 | - |
| 4 | 74 | 0.137 | -1/3 | - |
| 5 | 76 | 0.100 | -1/3 | MIM |
| 6 | 392 | 0.099 | -1/3 | - |
| 7 | 280 | 0.0864 | -1/3 | - |

For cancer networks, the three most frequently found 3-node motifs are id_6, id_12 and id_36. By examining the top three motifs, we observe the following common features: (i) they do not contain any subgraphs (*irreducible*); (ii) they are composed of a minimal number of edges (*N*-1 edges for a *N*-node motif); (iii) the reciprocity *r* values are negative (−0.5) and those motifs have *no* cycles; (iv) they account for at least 8% of the total number of motifs; and (v) they are associated with the lowest or the second lowest graph energies, regardless of the graph energy definition.

The motifs ranked 4^th^ to 7^th^ (“id_38,” “id_74,” “id_14,” and “id_98”) are composed of three edges. Motif “id_38” is the so-called feed-forward loop (FFL), which does not contain cycles, whereas both “id_74” and “id_14” contain cycles. The 7^th^-ranked motif (“id_98”) is the so-called 3-cycle. Motif “id_12” is a subgraph of “id_38,” “id_74,” and “id_98”, while “id_36” (MIM) is a subgraph of FFL and SIM is a subgraph of “id_14”. In other words, the frequently found motifs are the simplest motifs, and are subgraphs of more complex motifs.

For 4-node motifs, the above features (i) to (iv) (but not feature (v)) are also valid for the top seven most frequently found motifs. It is interesting to note that the *irreducible* and negative reciprocity value (that is, −1/3) features hold at the 4-node level. In addition, feature (v) holds if we consider graph energies *E, LE*, and *QE*, but not the other six graph energy definitions. Furthermore, the above five features also hold for STNs and cellular processes (see Supplementary File 1 - Supplementary Tables S9 and S10).

The ranges of average probability for the top three frequently found 3-node motifs and the top seven frequently found 4-node motifs are shown in Supplementary File 1 - Supplementary Tables S11. Other than the 4-node motifs of cellular processes, the ranges of average probability are quite similar. We note that cellular processes may differ from the other two families of networks, but it is not clear whether this is because a relatively small number (nine) of networks is available.

In Table 6, we summarize the top three most frequently found motifs and the top seven motifs identified among different networks. Cancer networks and STNs exhibit very similar results, which suggests that the underlying architectures are highly similar. This indicates that molecular networks are composed of a *finite* number of motif patterns—around seven patterns—with an upper *graph energy limit*. We conjecture that other molecular networks, such as cell cycles, may demonstrate similar properties.

**Table 6.** Comparison of frequently found motifs identified in cancer networks, STNs, and cellular processes.

|  | Cancer networks | STN | Cellular processes |
| --- | --- | --- | --- |
| Top three most frequently found motifs |  |  |  |
| 3-node ID | 6, 12, 36 | 12, 6, 36 | 12, 36, 6 |
| Rank of<br>*KC* | 1, 3, 2 | 3, 1, 2 | 3, 2, 1 |
| Top seven most frequently found motifs |  |  |  |
| 4-node ID | 14, 328, 28, 74, 76, 392, 280 | 14, 28, 74, 328, 280, 76, 392 | 392, 328, 76, 280, 2184, 74, 28 |
| Rank of<br>*KC* | 1, 7, 3, 5, 6, 33, 24 | 1, 3, 5, 24, 7, 24, 6, 33 | 33, 7, 6, 24, 7, 16, 5, 3 |

Next, we examine the association of frequently found motifs and complexity measures. From Table 6, we observe that frequently found motifs have a lower *KC* ranking (smaller *KC* value). A smaller *KC* value implies a lower degree of randomness, less information, and higher compressibility. However, this observation is not exactly the same at the 4-node level: there are three instances where the rank of *KC* is larger. For instance, the rank of *KC* is as high as 16, 24, and 33 for id_2184, id_280, and id_392, respectively. No obvious association exists between frequently found motifs and cyclomatic complexity measures.

In summary, according to the above findings, we hypothesize that there are underlying organizing principles that lead to the emergence of network structures.

### Network motifs absent from the network

We enumerated all possible 3-node motifs and 4-node motifs for the 17 cancer networks, 46 STNs, and nine cellular processes. In Supplementary File 1 - Supplementary Tables S12 summarizes the number of 3-node motifs and 4-node motifs with a non-zero frequency of occurrence in cancer networks, STNs, and cellular processes. It is interesting to see that certain motifs are never present in the three types of networks, except in two of the cellular processes. The first one is the “adherens junction” network which consist of a 3-node motif (id_110), composed of three genes: *CTNNA1, ACTB*, and *AFDN*. The second one is the “Signaling pathways regulating pluripotency of stem cells (hsa04550)” network. We have identified a fully connected 3-node motif (id_238) with three feedback loops connecting three genes: *Oct4, Sox2*, and *Nanog*. It is well-known that, with the *LIN28* genes, these four genes are liable to reprogram human somatic cells into pluripotent stem cells [68].

As long as there are motifs absent in the cancer networks, STNs, and cellular processes, there is a graph energy cutoff associated with these three families of networks. Supplementary File 1 - Supplementary Tables S13 depicts the results of the energy cutoffs, maximum graph energies, and their ratios for the cancer networks. Among the nine graph energy definitions, the ratios may be as high as 0.750 and 0.667 for 3-node motifs and 4-node motifs, respectively. However, they can be as low as 0.536 (*LQ*^*t*^ energy) and 0.481 (*AQ*^*t*^ energy) for the 3-node motifs and 4-node motifs, respectively. The results of the graph energy cutoffs and ratios for the STNs and cellular processes are given in Supplementary File 1 - Supplementary Tables S14 and S15. For cellular processes, the cutoff ratio may be as high as 1.00 for the 3-node motifs, because we have identified a fully connected 3-node motif (id_238). At the 3-node level, two of the cellular processes exhibit peculiar network structures; this is an open issue that remains to be investigated.

Nevertheless, our results suggest that there is an existing energy cutoff or ratio that constrains the presence of certain motifs embedded in a molecular network. In addition, the data indicate that the cutoff ratio for 3-node motifs is slightly higher than that for 4-node motifs. Furthermore, the motif probability distribution density and the graph energy of a motif obeys an inverse relation, that is, the smaller the probability, the higher the graph energy.

### Characterizing the frequency distributions of motifs using entropy

We utilized the entropy-based quantity, normalized Shannon entropy, *H*_*R*_, to quantify the frequency distributions of the occurrence of motifs for the cancer networks. For randomized distribution, *H* achieves the maximal values, 3.700 (log_2_ (13)) and 7.637 (log_2_ (199)), for 3-node motifs and 4-node motifs, respectively. Table 7 lists the number of 3-node motifs, *N*_*3*_; normalized Shannon entropy, *H*_*3R*_, for the 3-node motifs; number of 4-node motifs, *N*_*4*_; and normalized Shannon entropy, *H*_*4R*_, for the 4-node motifs.

**Table 7.** The results of *N*_*3*_, *N*_*4*_, *H*_*3R*_ and *H*_*4R*_ for cancer networks.

| Cancer networks | $N_3$ | $H_{3R}$ | $N_4$ | $H_{4R}$ |
| --- | --- | --- | --- | --- |
| Acute_myeloid_leukemia_[hsa05221] | 160 | 0.428 | 577 | 0.385 |
| Basal_cell_carcinoma_[hsa05217] | 34 | **0.323** | 51 | **0.273** |
| Breast_cancer_[hsa05224] | 145 | 0.449 | 445 | 0.437 |
| Choline_metabolism_in_cancer_[hsa05231] | 70 | 0.465 | 193 | 0.406 |
| Chronic_myeloid_leukemia_[hsa05220] | 71 | 0.368 | 145 | 0.323 |
| Colorectal_cancer_[hsa05210] | 71 | 0.467 | 124 | 0.403 |
| Endometrial_cancer_[hsa05213] | 45 | 0.329 | 60 | 0.295 |
| Gastric_cancer_[hsa05226] | 87 | 0.355 | 153 | 0.344 |
| Glioma_[hsa05214] | 80 | 0.410 | 183 | 0.390 |
| Hepatocellular_carcinoma_[hsa05225] | 65 | 0.355 | 80 | 0.350 |
| Melanoma_[hsa05218] | 46 | 0.374 | 82 | 0.337 |
| Non-small_cell_lung_cancer_[hsa05223] | 103 | **0.493** | 284 | **0.483** |
| Pancreatic_cancer_[hsa05212] | 74 | 0.422 | 131 | 0.371 |
| Pathways_in_cancer_[hsa05200] | 640 | 0.473 | 2795 | 0.450 |
| Prostate_cancer_[hsa05215] | 102 | 0.372 | 357 | 0.335 |
| Renal_cell_carcinoma_[hsa05211] | 58 | 0.385 | 114 | 0.353 |
| Small_cell_lung_cancer_[hsa05222] | 61 | 0.362 | 96 | 0.318 |

For all of the cancer networks we studied, the frequency distributions of the motifs are not uniformly distributed among the motif patterns; therefore, *H*_*3R*_ and *H*_*4R*_ are different from one another. The results of *N*_*3*_, *N*_*4*_ *H*_*3R*_, and *H*_*4R*_ for STNs and cellular processes are given in Supplementary File 1 - Supplementary Tables S16 and S17, respectively.

Supplementary File 1 - Supplementary Table S18 shows the ranges of *H*_*3R*_ and *H*_*4R*_ for cancer networks, STNs, and cellular processes. The ranges of *H*_*3R*_ and *H*_*4R*_ for cancer networks and cellular processes are quite similar, whereas STNs show larger ranges: 0.532 and 0.622. We also note that the HIF-1 signaling pathway [hsa04066] has very small *H*_*3R*_ and *H*_*4R*_. This is because the transcription factor, HIF-1β, functions as a master regulator of many genes; therefore, the SIM motif is the dominant motif at both 3-node and 4-node levels.

## Discussion and conclusions

Network motifs play an important role in biological networks. We made use of a rigorous mathematical and systematic approach—spectral graph theory—to characterize 3-node and 4-node network motifs. A total of nine graph energies were introduced to characterize network motifs. In addition, we characterized their complexity by using two widely accepted complexity measures, *CC* and *KC*.

The concept of a unique identifier was introduced to label network motifs. This novel idea combines four parameters—graph energy, reciprocity, *CC*, and eigenvalue spectrum—to characterize a network motif.

A foreseeable application of this identifier is to examine the transition between different motifs. It is possible that the regulatory interactions among the genetic elements may be disrupted or activated because of genetic mutation or epigenetic modification.

We conjecture that driver mutations are enriched or depleted in certain motif positions, such as the source node of a motif. A source node is a node that has only outgoing edges. In other words, a mutation driver gene acts as an upstreaming regulator. Several studies have reported that certain motif positions, such as the source nodes and target nodes, are enriched in cancer-associated genes [69-70].

Some of the 3-node motifs and 4-node motifs are interconnected through shared genetic elements. These types of modules are called coupled motif structures (CMS) in our previous work [45]. One can merge interconnected motifs to form higher-order network structures.

We have extended a developed algorithm to identify complete sets of 3-node motifs and 4-node motifs for 17 cancer networks, 46 STNs, and nine cellular processes. Except for a few networks, 3-node motifs and 4-node motifs account for more than 70% of the nodes in the studied networks. Furthermore, this study discovered the following features:

(i). The relative entropies of the motif distributions are not equal, or close to, one, indicating that the identified motifs are not distributed uniformly among the 13 and 199 patterns.

(ii) Biological networks are composed of a finite number of motif patterns, this suggest the presence of graph energy cutoffs.

(iii) Irreducible motif patterns are the most frequently found motifs; for instance, the cascade pattern is the most frequently found motif, followed by the SIM and MIM motifs.

(iv) All of the three families of networks exhibit the above features, suggesting that there is a universal organization principle determining the underlying network architecture.

In conclusion, this study provides a systematic and rigorous approach to dissecting the underlying structures of biological molecular networks. SGT serves as a powerful approach in distinguishing different motif topologies or connectivity patterns, which inspires our hypothesis that network structures can be understood in terms of the 3-node and 4-node motifs. The next step is to test our hypothesis by analyzing the 5-node motifs. We expect that our efforts may help to elucidate the complex nature of molecular networks.

## Supporting information

Supplementary File 1. Input data and Methods

Supplementary Table S1. List of the nodes, edges, and motif-associated nodes information for the 17 cancer networks.

Supplementary Table S2. List of the nodes, edges, and motif-associated nodes information for the 46 STNs.

Supplementary Table S3. List of the nodes, edges, and motif-associated nodes information for the nine cellular processes.

Supplementary Table S4. Motif identification tool - *PatternFinder* algorithm.

Supplementary Table S5. Algorithm for finding a unique identifier for the 3-node motifs.

Supplementary Table S6. The results of the lower (*E*_*min*_) and upper (*E*_*max*_) bounds of the nine graph energies and ratios for the 3-node motifs and 4-node motifs.

Supplementary Table S7. The results of the correlation strength (*SRCC*) between graph complexity and graph energy for the 3-node motifs and 4-node motifs.

Supplementary Table S8. The results of the correlation strength (including minimum, maximum, and ranges) between graph complexity and graph energy for the 3-node motifs and 4-node motifs.

Supplementary Table S9. The top seven most frequently found 3-node motifs and 4-node motifs for STNs.

Supplementary Table S10. The top seven most frequently found 3-node motifs and 4-node motifs for cellular processes.

Supplementary Table S11. The ranges of average probability for the top three frequently found 3-node motifs and the top seven frequently found 4-node motifs.

Supplementary Table S12. The results of the number of possible 3-node motif patterns and 4-node motif patterns present in the 17 cancer networks, 46 STNs, and nine cellular processes.

Supplementary Table S13. The results of the cutoff and maximum graph energies of the smallest non-zero average probability for the 3-node motifs and 4-node motifs present in cancer networks.

Supplementary Table S14. The results of the cutoff and maximum graph energies of the smallest non-zero average probability for the 3-node motifs and 4-node motifs present in STNs.

Supplementary Table S15. The results of the cutoff and maximum graph energies of the smallest non-zero average probability for the 3-node motifs and 4-node motifs present in cellular processes.

Supplementary Table S16. The results of *N*_*3*_, *N*_*4*_, *H*_*3R*_, and *H*_*4R*_ for STNs.

Supplementary Table S17. The results of *N*_*3*_, *N*_*4*_, *H*_*3R*_, and *H*_*4R*_ for cellular processes.

Supplementary Tables S18. The ranges of *H*_*3R*_ and *H*_*4R*_ for cancer networks, STNs, and cellular processes.

Supplementary File 2. The results of both 3-node and 4-node motif subgraphs. Supplementary File 3. The results of the 4-node motif graph energies and eigenvectors.

Supplementary File 4. The complete sets of reciprocity values of the 4-node motifs.

Supplementary File 5. The complete sets of cyclomatic complexity and Kolmogorov complexity values of the 4-node motifs.

### Acknowledgments

Dr. Chien-Hung Huang and Mr. Alice, Hsieh work are supported by the Ministry of Science and Technology (MOST, https://www.most.gov.tw/?l=en) under the grants of MOST 107-2221-E-150-038. Dr. Ka-Lok Ng work is supported by the grants of MOST 106-2221-E-468-017, MOST 107-2632-E-468-002, and also supported under the grants from Asia University, 106-asia-06, 107-asia-02 and 107-asia-09. Dr. Jeffrey J. P. Tsai work is supported by the grant of MOST 107-2632-E-468-002. We would like to thank He-Xing Li, Ci-Jun Peng and I-Lun Hsieh, who spent efforts on developing the codes. The funders, MOST and Asia University, had no role in study design, data collection and analysis, decision to publish, or preparation of the manuscript. We also thank the ‘Editage Professional English Editing Service, Cactus Communications’, for editing the English.

## Authors’ Contributions

Chien-Hung Huang conducted the algorithm development, review and edited the manuscript. Jeffrey J. P. Tsai provided interpretation of the results and review the manuscript. Nilubon Kurubanjerdjit conducted the complexity measure analysis, literature search and participated in discussion. Ka-Lok Ng is the corresponding author, who designed the study, review and drafted the manuscript.

