## Supplementart file 1 for "Computational analysis of molecular networks using spectral graph theory, complexity measures and information theory"

### Supplementary File 1

### Dissecting the Organization of Molecular Networks Using Network motifs

Chien-Hung Huang[1], Jeffrey J. P. Tsai[2], Nilubon Kurubanjerdjit[3]
Ka-Lok Ng[2,4] *

[1]*Department of Computer Science and Information Engineering, National Formosa University, Yun-Lin, Taiwan*

[2]*Department of Bioinformatics and Medical Engineering, Asia University, Taichung, Taiwan*

[3]*SIQ-DIT research group, School of Information Technology, Mae Fah Luang University, Chiang Rai, Thailand*

[4]*Department of Medical Research, China Medical University Hospital, China Medical University, Taichung, Taiwan*

[1]** [2] , [2]** [3]**

** corresponding author*

[2,4 *]**

#### Methods

Supplementary File 1 – Supplementary Table S1 summarizes the nodes, edges and motif-associated node information for the 17 cancer networks. Besides the 'Hepatocellular carcinoma' and 'Prostate cancer' networks, the percentages of nodes associated with motifs are more than 70%. The results suggest that genes embedded in the 3-node motifs and 4-node motifs account for the major portion of the cancer networks.

We noticed that a few of the networks' KGML files have missing information. Feedback loop information is not recorded in the KGML, but it is displayed on the KEGG webpage. For instance, the 'Signaling pathways regulating pluripotency of stem cells' pathway consists of the feedback regulatory relations among three genes, *Nanog*, *Oct4* and *Sox2*. The KGML file provides the regulatory relations among *Oct4* → *Nanog* → *Sox2* but not *Nanog* → *Oct4* → *Sox2*. To remedy this problem, we have to examine each analyzed network manually, and insert the missing information back in the KGML files.

Supplementary Table S1. List of the nodes, edges and motif-associated nodes information for the 17 cancer networks.

| Cancer networks | nodes | edges | motif-associated nodes |
|---|---|---|---|
| Acute_myeloid_leukemia_[hsa05221] | 43 | 53 | 100% |
| Basal_cell_carcinoma_[hsa05217] | 23 | 20 | 91.3% |
| Breast_cancer_[hsa05224] | 63 | 60 | 81.0% |
| Choline_metabolism_in_cancer_[hsa05231] | 37 | 34 | 73.0% |
| Chronic_myeloid_leukemia_[hsa05220] | 47 | 39 | 78.7% |
| Colorectal_cancer_[hsa05210] | 48 | 39 | 77.1% |
| Endometrial_cancer_[hsa05213] | 37 | 27 | 78.4% |

| | | | |
|---|---|---|---|
| Gastric_cancer_[hsa05226] | 73 | 51 | 71.2% |
| Glioma_[hsa05214] | 36 | 36 | 86.1% |
| Hepatocellular_carcinoma_[hsa05225] | 82 | 49 | 64.6% |
| Melanoma_[hsa05218] | 28 | 25 | 89.3% |
| Non-small_cell_lung_cancer_[hsa05223] | 43 | 43 | 83.7% |
| Pancreatic_cancer_[hsa05212] | 50 | 46 | 92.0% |
| Pathways_in_cancer_[hsa05200] | 152 | 197 | 94.7% |
| Prostate_cancer_[hsa05215] | 53 | 38 | 67.9% |
| Renal_cell_carcinoma_[hsa05211] | 40 | 27 | 72.5% |
| Small_cell_lung_cancer_[hsa05222] | 48 | 34 | 75.0% |

Supplementary Table S2. List of nodes, edges and motif-associated nodes information for the 46 STN.

| STN | nodes | edges | motif-associated nodes |
|---|---|---|---|
| Adipocytokine_signaling_pathway_[hsa04920] | 36 | 43 | 91.7% |
| AMPK_signaling_pathway_[hsa04152] | 61 | 52 | 75.4% |
| Apelin_signaling_pathway_[hsa04371] | 55 | 56 | 85.5% |
| B_cell_receptor_signaling_pathway_[hsa04662] | 46 | 44 | 73.9% |
| Calcium_signaling_pathway_[hsa04020] | 44 | 28 | 54.5% |
| cAMP_signaling_pathway_[hsa04024] | 74 | 71 | 90.5% |
| cGMP-PKG_signaling_pathway_[hsa04022] | 59 | 52 | 83.1% |
| Chemokine_signaling_pathway_[hsa04062] | 51 | 58 | 94.1% |
| C-type_lectin_receptor_signaling_pathway_[hsa04625] | 75 | 90 | 96.0% |
| ErbB_signaling_pathway_[hsa04012] | 53 | 82 | 98.1% |
| Estrogen_signaling_pathway_[hsa04915] | 34 | 33 | 88.2% |
| Fc_epsilon_RI_signaling_pathway_[hsa04664] | 40 | 35 | 75.0% |
| FoxO_signaling_pathway_[hsa04068] | 74 | 69 | 91.9% |
| Glucagon_signaling_pathway_[hsa04922] | 45 | 38 | 68.9% |
| GnRH_signaling_pathway_[hsa04912] | 40 | 38 | 90.0% |
| Hedgehog_signaling_pathway_[hsa04340] | 23 | 35 | 100.0% |
| HIF-1_signaling_pathway_[hsa04066] | 62 | 53 | 83.9% |
| Hippo_signaling_pathway_[hsa04390] | 78 | 60 | 76.9% |
| Insulin_signaling_pathway_[hsa04910] | 63 | 69 | 87.3% |
| Jak-STAT_signaling_pathway_[hsa04630] | 34 | 34 | 97.1% |
| MAPK_signaling_pathway_[hsa04010] | 115 | 163 | 98.3% |
| mTOR_signaling_pathway_[hsa04150] | 67 | 72 | 83.6% |
| Neurotrophin_signaling_pathway_[hsa04722] | 74 | 112 | 95.9% |
| NF-kappa_B_signaling_pathway_[hsa04064] | 101 | 76 | 69.3% |
| NOD-like_receptor_signaling_pathway_[hsa04621] | 114 | 131 | 84.2% |
| Notch_signaling_pathway_[hsa04330] | 27 | 16 | 63.0% |
| Oxytocin_signaling_pathway_[hsa04921] | 52 | 56 | 88.5% |
| p53_signaling_pathway_[hsa04115] | 59 | 57 | 88.1% |
| Phosphatidylinositol_signaling_system_[hsa04070] | 30 | 60 | 90.0% |
| Phospholipase_D_signaling_pathway_[hsa04072] | 51 | 45 | 84.3% |
| PI3K-Akt_signaling_pathway_[hsa04151] | 90 | 77 | 76.7% |
| PPAR_signaling_pathway_[hsa03320] | 51 | 55 | 88.2% |
| Prolactin_signaling_pathway_[hsa04917] | 39 | 44 | 97.4% |
| Rap1_signaling_pathway_[hsa04015] | 81 | 88 | 81.5% |
| Ras_signaling_pathway_[hsa04014] | 68 | 96 | 95.6% |
| Relaxin_signaling_pathway_[hsa04926] | 55 | 68 | 89.1% |
| RIG-I-like_receptor_signaling_pathway_[hsa04622] | 56 | 40 | 58.9% |
| Sphingolipid_signaling_pathway_[hsa04071] | 55 | 51 | 81.8% |
| T_cell_receptor_signaling_pathway_[hsa04660] | 67 | 67 | 76.1% |

| | | | |
|---|---|---|---|
| TGF-beta_signaling_pathway_[hsa04350] | 50 | 38 | 74.0% |
| Thyroid_hormone_signaling_pathway_[hsa04919] | 65 | 56 | 75.4% |
| TNF_signaling_pathway_[hsa04668] | 77 | 48 | 54.5% |
| Toll_and_Imd_signaling_pathway_[dme04624] | 55 | 59 | 92.7% |
| Toll-like_receptor_signaling_pathway_[hsa04620] | 73 | 91 | 90.4% |
| VEGF_signaling_pathway_[hsa04370] | 28 | 32 | 100.0% |
| Wnt_signaling_pathway_[hsa04310] | 69 | 84 | 95.7% |

Supplementary Table S3. List of the nodes, edges and motif-associated nodes information for the nine cellular processes.

| Cell cycle | nodes | edges | motif-associated nodes |
|---|---|---|---|
| Adherens junction_[hsa04520] | 83 | 80 | 84.6% |
| Apoptosis [hsa04210] | 123 | 132 | 89.8% |
| Cell_cycle [hsa04110] | 115 | 79 | 61.1% |
| Cellular senescence [hsa04218] | 122 | 104 | 78.2% |
| Focal adhesion [hsa04510] | 72 | 101 | 98.4% |
| Gap junction [hsa04540] | 59 | 56 | 100.0% |
| Necroptosis [hsa04217] | 94 | 77 | 89.4% |
| Regulation of actin cytoskeleton [hsa04810] | 86 | 75 | 82.4% |
| Signaling pathways regulating pluripotency of stem cells [hsa04550] | 108 | 64 | 58.2% |

Supplementary Table S4. Motif identification tool - *PatternFinder* algorithm

There are many network motif detection tools, such as, FANMOD, MAVISTO, MFINDER, NetMatch and SNAVI. As we have reported that those tools have a few limitations[1] on motif identification: (i) the subgraphs may not be recoverable due to the use of randomize algorithm, and (ii) subgraph's nodes identities were missing.

An algorithm named *PatternFinder* was developed to identify: (i) functional motifs embedded in the 3-node motifs and 4-node motifs, and (ii) motifs compose of three nodes and four nodes in a network[2]. The advantage of *PatternFinder* is its ability to identify subgraphs that are not identified by MFINDER. As shown in Supplementary Figure 1, given the "Input network", *PatternFinder* is able to identify two four-node motifs, i.e. motif 'id_904' and motif 'id_906', whereas MFINDER can identify motif 'id_906' only. It is because MFINDER recognizes motif 'id_904' is a subgraph of motif 'id_906'. In other words, *PatternFinder* is able to identify subgraphs embedded in a motif.

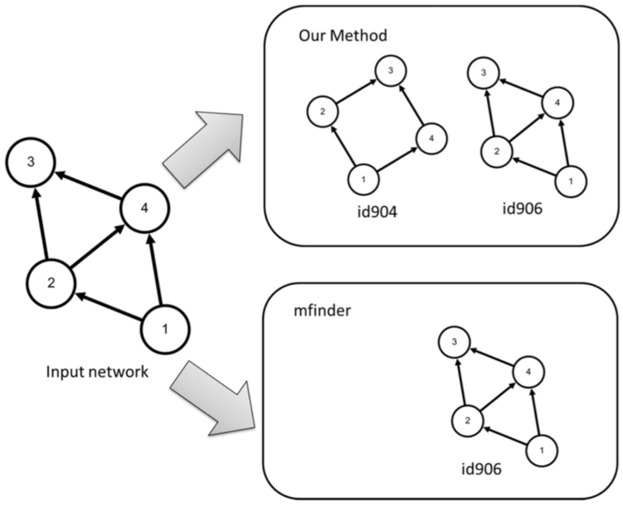


Supplementary Figure 1 A comparison of two motif identification algorithms: *mfinder* and *PatternFinder*

In the following, a 4-node motif is used as an example to illustrate the basic concept behind the *PatternFinder* algorithm. Given a network called '*net'* with 20 nodes, an adjacency matrix can be constructed. Let *n* denotes the total number of nodes. Assuming that we want to identify a motif which is denoted by the integer '2204', *PatternFinder* read in the '2204' pattern. This motif composes of four nodes and five edges, where the edges are denoted by $\boldsymbol{t_0}$, $\boldsymbol{t_1}$, $\boldsymbol{t_2}$, $\boldsymbol{tr_0}$ and $\boldsymbol{tr_1}$. Starting from node A, *PatternFinder* begins to examine the following patterns: (i) is node A and node B connects with an edge $\boldsymbol{t_0}$, (ii) is node B and node C connects with an edge $\boldsymbol{t_1}$, (iii) is node C and node D connects with an edge $\boldsymbol{t_2}$, and (iv) is node D and node A connects with an edge $\mathrm{t}_0$, and node D and node B connects with an edge $\boldsymbol{tr_1}$.

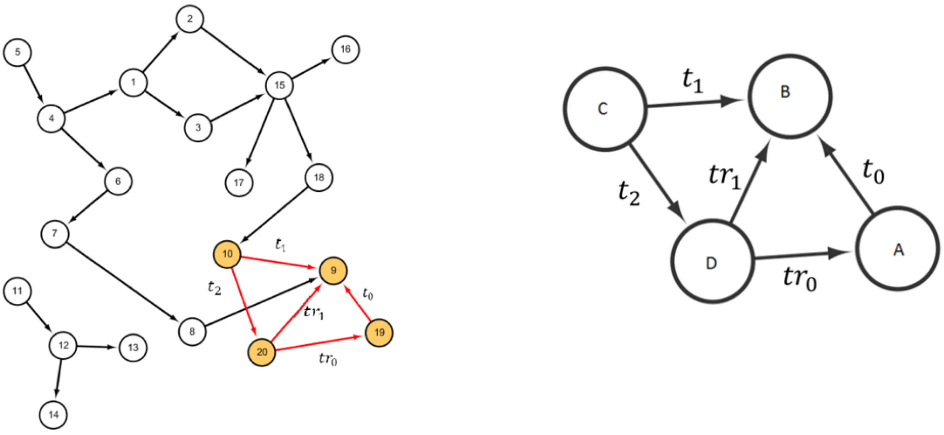


Supplementary Figure 2 (a) an input network named '*net'* and (b) the 4-node motif 'id_2204'

Starting from the network named '*net*', the algorithm begins the search from node 1 and labels it as node A. Node 1 and node 2 or node B are linked, the edge is denoted by *edge*(1, 2). The algorithm continues to search if there is a node links to node B, if not, the algorithm will relabel node B to node 3 and repeat the search. From Supplementary Figure 2, it was found that A=19, B=9, C=10, and D=20 are connected by three edges, i.e. *edge*(19,9)= $\boldsymbol{t_0}$ , *edge*(9,10)= $\boldsymbol{t_1}$, *edge*(10,20)= $\boldsymbol{t_2}$ , hence, four nodes are identified. However, according to the 'id_2204' motif, there are two more edges need to be determined, i.e. *edge*(20,9)= $\boldsymbol{tr_1}$ and *edge*(20,19)= $\boldsymbol{tr_2}$ . The computational time complexity of the algorithm *PatternFinder* is $O(n^4)$.


Supplementary Table S5. Algorithm for finding a unique identifier for the 3-node motifs

The following algorithm is used to derive the subsets of parameters (energy parameters in *ES, r* and *CC*), which could separate the most number of motif types. In this algorithm, *ES* denotes the set of the graph energy parameters.

Input:

*ES*={$E_1$, $E_2$, … , $E_6$}, parameter *r*, parameter *CC* and all of the 13 motif types of the 3-node motif;

Output:

Fingerprint parameter set *MS* for the 3-node motif;

Step 1: For each parameter (energy parameter in *ES*, parameters *r* or *CC*), generate 13 sets corresponding to the 13 motif types; that is, one set corresponds to one motif type;

Step 2: For each parameter, calculate the corresponding values for all of the 13 motif types, and merge the corresponding sets generated in Step 1 into one set if they have the same computed value (this means the corresponding motif types cannot be distinguished by this parameter);

Step 3: *MS*={*CC*};

Step 4: For the current sets of parameter *CC*, if any set has more than one element (motif type) and some of the corresponding motif types can be distinguished by parameter *r*, then divide the set into the corresponding disjoint sets;

Step 5: $MS=MS \cup \{r\}$;

Step 6: Find the element in *ES* called $E_i$, which can separate the highest number of sets for the current sets of parameter *CC*; return $MS=MS \cup \{E_i\}$;

Supplementary Table S6. The results of the lower ($E_{min}$) and upper ($E_{max}$) bounds of the nine graph energies and ratios for the 3-node motifs and 4-node motifs.

| | | *E* | *LE* | *QE* | $AA^t$ | $LL^t$ | $QQ^t$ | $AL^t$ | $AQ^t$ | $LQ^t$ |
|---|---|---|---|---|---|---|---|---|---|---|
| 3-node | $E_{min}$ | 0.00 | 2.67 | 2.67 | 1.41 | 4.32 | 4.32 | 1.41 | 1.41 | 3.83 |
| | $E_{max}$ | 4.00 | 8.00 | 8.00 | 4.00 | 12.00 | 12.00 | 6.47 | 6.93 | 11.21 |
| | *ratio* | 2.00 * | 3.00 | 3.00 | 3.05 | 2.78 | 2.78 | 4.59 | 4.91 | 2.93 |
| 4-node | $E_{min}$ | 0.00 | 4.50 | 4.50 | 1.73 | 6.36 | 6.36 | 1.73 | 1.73 | 5.61 |
| | $E_{max}$ | 6 | 18 | 18 | 6 | 24 | 24 | 10.94 | 11.9 | 22.94 |
| | *ratio* | 3.00 | 4.00 | 4.00 | 3.47 | 3.77 | 3.77 | 6.32 | 6.88 | 4.09 |

The symbol '*' means $E_{min}$ is given by the second smallest graph energy *E*

Supplementary Table S7. The results of the correlation strength (*SRCC*) between graph complexity and graph energy for the 3-node motifs and 4-node motifs.

| | | *E* | *LE* | *QE* | *AA'* | *LL'* | *QQ'* | *AL'* | *AQ'* | *LQ'* |
|---|---|---|---|---|---|---|---|---|---|---|
| 3-node | *CC* | -0.003 | 0.224 | 0.171 | 0.032 | ***0.270***[*] | 0.188 | -0.023 | 0.062 | 0.257 |
| | *KC* | 0.484 | 0.685 | 0.698 | 0.617 | ***0.779***[*] | 0.748 | 0.359 | 0.604 | 0.773 |
| 4-node | *CC* | 0.332 | 0.630 | 0.614 | 0.453 | ***0.686***[*] | 0.651 | 0.384 | 0.452 | 0.670 |
| | *KC* | 0.602 | 0.764 | 0.766 | 0.712 | 0.771 | 0.782 | 0.648 | 0.684 | ***0.773***[*] |

[*] Bold-faced and italic numbers denote the maximum *SRCC* for 3-node motifs and 4-node motifs among the nine energies.

Supplementary Table S8. The results of the correlation strength (including minimum, maximum and ranges) between graph complexity and graph energy for the 3-node motifs and 4-node motifs.

| | | minimum | maximum | range |
|---|---|---|---|---|
| 3-node | (*CC*, energy) | -0.023 | 0.270 | 0.293 |
| | (*KC*, energy) | 0.359 | 0.779 | 0.420 |
| 4-node | (*CC*, energy) | 0.332 | 0.686 | 0.354 |
| | (*KC*, energy) | 0.602 | 0.782 | 0.180 |

Supplementary Table S9. The top seven most frequently found 3-node motifs and 4-node motifs for STN.

| rank | ID | average probability | *r* | Name & embedded motif ID |
|---|---|---|---|---|
| 3-node | | | | |
| 1 | 12 | 0.402 | -1/2 | Cascade |
| 2 | 6 | 0.380 | -1/2 | SIM |
| 3 | 36 | 0.194 | -1/2 | MIM |
| 4 | 38 | 0.0154 | -1 | FFL, 12, 36 |
| 5 | 14 | 0.00360 | 1/3 | 6 |
| 6 | 74 | 0.00250 | 1/3 | 12, 36 |
| 7 | 98 | 0.00138 | -1 | 12 |
| 4-node | | | | |
| 1 | 14 | 0.260 | -1/3 | SIM |
| 2 | 28 | 0.150 | -1/3 | - |
| 3 | 74 | 0.127 | -1/3 | null |
| 4 | 328 | 0.112 | -1/3 | Cascade |
| 5 | 280 | 0.103 | -1/3 | - |
| 6 | 76 | 0.0932 | -1/3 | MIM |
| 7 | 392 | 0.0908 | -1/3 | - |

Supplementary Table S10. The top seven most frequently found 3-node motifs and 4-node motifs for cellular processes.

| rank | ID | average probability | *r* | Name & embedded motif ID |
|---|---|---|---|---|
| 3-node | | | | |
| 1 | 12 | 0.443 | -1/2 | Cascade |
| 2 | 36 | 0.316 | -1/2 | MIM |
| 3 | 6 | 0.183 | -1/2 | SIM |
| 4 | 38 | 0.0206 | -1 | FFL, 12, 36 |
| 5 | 74 | 0.0167 | 1/3 | 12, 36 |
| 6 | 14 | 0.00790 | 1/3 | 6 |
| 7 | 46 | 0.00250 | -1/2 | 6, 12, 14, 36, 38 |
| 4-node | | | | |
| 1 | 392 | 0.159 | -1/3 | - |
| 2 | 328 | 0.145 | -1/3 | Cascade |
| 3 | 76 | 0.121 | -1/3 | MIM |
| 4 | 280 | 0.120 | -1/3 | - |
| 5 | 2184 | 0.115 | -1/3 | MIM |
| 6 | 74 | 0.0990 | -1/3 | - |
| 7 | 28 | 0.0861 | -1/3 | - |

Supplementary Tables S11. The ranges of average probability for the top three 3-node motifs and the top seven 4-node motifs.

| Network type | 3-node motifs | 4-node motifs |
|---|---|---|
| Cancer networks | 0.152 ~ 0.421 = 0.269 | 0.0864 ~ 0.224 = 0.138 |
| STN | 0.194 ~0.402 = 0.208 | 0.0908 ~ 0.260 = 0.169 |
| cellular processes | 0.183 ~ 0.443 = 0.260 | 0.0861 ~ 0.159 = 0.073 |

Supplementary Tables S12. The results of the number of possible 3-node motifs patterns and 4-node motifs patterns present in the 17 cancer networks, 46 STN and nine cellular processes.

| | cancer network | STN | cellular process |
|---|---|---|---|
| 3-node | 7/13 | 11/13 | 13/13 |
| 4-node | 38/199 | 88/199 | 77/199 |

Supplementary Tables S13. The results of the cutoff and maximum graph energies of the smallest non-zero average probability for the 3-node motif and 4-node motif present in cancer networks.

| | energy | *E* | *LE* | *QE* | *AA'* | *LL'* | *QQ'* | *AL'* | *AQ'* | *LQ'* |
|---|---|---|---|---|---|---|---|---|---|---|
| 3-node | cutoff | 3 | 4.292 | 4.464 | 3 | 6.292 | 6.464 | 4.252 | 4.363 | 6.012 |
| | max | 4 | 8 | 8 | 4 | 12 | 12 | 6.472 | 6.928 | 11.21 |
| | ratio | 0.750 | 0.537 | 0.558 | 0.750 | 0.524 | 0.539 | 0.657 | 0.630 | 0.536 |
| 4-node | cutoff | 4.00 | 9.00 | 9.00 | 4.00 | 12.54 | 12.44 | 5.723 | 5.723 | 12 |
| | max | 6 | 18 | 18 | 6 | 24 | 24 | 10.94 | 11.9 | 22.94 |
| | ratio | 0.667 | 0.500 | 0.500 | 0.667 | 0.523 | 0.518 | 0.523 | 0.481 | 0.523 |

ratio = cutoff / max

Supplementary Table S14. The results of the cutoff and maximum graph energies of the smallest non-zero average probability for the 3-node motif and 4-node motif present in STN.

| | | $E$ | $LE$ | $QE$ | $AA'$ | $LL'$ | $QQ'$ | $AL'$ | $AQ'$ | $LQ'$ |
|---|---|---|---|---|---|---|---|---|---|---|
| 3-node | cutoff | 3.062 | 5.333 | 5.56 | 3.236 | 8.243 | 8.247 | 4.762 | 5.022 | 7.586 |
| | max | 4 | 8 | 8 | 4 | 12 | 12 | 6.472 | 6.928 | 11.21 |
| | ratio | 0.766 | 0.667 | 0.695 | 0.809 | 0.687 | 0.687 | 0.736 | 0.725 | 0.677 |
| 4-node | cutoff | 4.395 | 10.58 | 10.64 | 4.705 | 14.23 | 14.23 | 7.453 | 7.73 | 13.68 |
| | max | 6 | 18 | 18 | 6 | 24 | 24 | 10.94 | 11.9 | 22.94 |
| | ratio | 0.733 | 0.588 | 0.591 | 0.784 | 0.593 | 0.593 | 0.681 | 0.650 | 0.596 |

ratio = cutoff / max

.

Supplementary Table S15. The results of the cutoff and maximum graph energies of the smallest non-zero average probability for the 3-node motif and 4-node motif present in cellular processes.

| | | $E$ | $LE$ | $QE$ | $AA'$ | $LL'$ | $QQ'$ | $AL'$ | $AQ'$ | $LQ'$ |
|---|---|---|---|---|---|---|---|---|---|---|
| 3-node | cutoff | 4 | 8 | 8 | 4 | 12 | 12 | 6.472 | 6.928 | 11.21 |
| | max | 4 | 8 | 8 | 4 | 12 | 12 | 6.472 | 6.928 | 11.21 |
| | ratio | 1 | 1 | 1 | 1 | 1 | 1 | 1 | 1 | 1 |
| 4-node | cutoff | 4.116 | 10.5 | 10.5 | 4.36 | 14.26 | 14.21 | 6.815 | 7.07 | 13.28 |
| | max | 6 | 18 | 18 | 6 | 24 | 24 | 10.94 | 11.9 | 22.94 |
| | ratio | 0.686 | 0.583 | 0.583 | 0.727 | 0.594 | 0.592 | 0.623 | 0.594 | 0.579 |

ratio = cutoff / max

.

Supplementary Table S16. The results of $N_3$, $N_4$, $H_{3R}$ and $H_{4R}$ for STN.

| STN | $N_3$ | $H_{3R}$ | $N_4$ | $H_{4R}$ |
|---|---|---|---|---|
| Adipocytokine_signaling_pathway_[hsa04920] | 118 | 0.426 | 386 | 0.414 |
| AMPK_signaling_pathway_[hsa04152] | 262 | 0.345 | 1830 | 0.308 |
| Apelin_signaling_pathway_[hsa04371] | 127 | 0.372 | 358 | 0.351 |
| B_cell_receptor_signaling_pathway_[hsa04662] | 96 | 0.419 | 311 | 0.411 |
| Calcium_signaling_pathway_[hsa04020] | 60 | 0.406 | 124 | 0.382 |
| cAMP_signaling_pathway_[hsa04024] | 353 | 0.217 | 2289 | 0.142 |
| cGMP-PKG_signaling_pathway_[hsa04022] | 202 | 0.290 | 1002 | 0.209 |
| Chemokine_signaling_pathway_[hsa04062] | 144 | 0.412 | 508 | 0.391 |
| C-type_lectin_receptor_signaling_pathway_[hsa04625] | 239 | 0.486 | 716 | 0.449 |
| ErbB_signaling_pathway_[hsa04012] | 375 | 0.442 | 2054 | 0.433 |
| Estrogen_signaling_pathway_[hsa04915] | 49 | 0.365 | 97 | 0.332 |
| Fc_epsilon_RI_signaling_pathway_[hsa04664] | 87 | 0.451 | 321 | 0.427 |
| FoxO_signaling_pathway_[hsa04068] | 902 | 0.389 | 12174 | 0.268 |
| Glucagon_signaling_pathway_[hsa04922] | 78 | 0.326 | 182 | 0.270 |
| GnRH_signaling_pathway_[hsa04912] | 76 | 0.323 | 212 | 0.315 |
| Hedgehog_signaling_pathway_[hsa04340] | 147 | 0.518 | 752 | 0.484 |
| HIF-1_signaling_pathway_[hsa04066] | 407 | **0.120** | 3382 | **0.034** |
| Hippo_signaling_pathway_[hsa04390] | 162 | 0.446 | 511 | 0.371 |
| Insulin_signaling_pathway_[hsa04910] | 188 | 0.452 | 659 | 0.431 |
| Jak-STAT_signaling_pathway_[hsa04630] | 179 | 0.398 | 894 | 0.318 |
| MAPK_signaling_pathway_[hsa04010] | 745 | 0.451 | 4280 | 0.436 |
| mTOR_signaling_pathway_[hsa04150] | 209 | 0.483 | 801 | 0.488 |
| Neurotrophin_signaling_pathway_[hsa04722] | 525 | 0.432 | 2957 | 0.427 |
| NF-kappa_B_signaling_pathway_[hsa04064] | 304 | 0.443 | 1586 | 0.325 |
| NOD-like_receptor_signaling_pathway_[hsa04621] | 566 | 0.454 | 2851 | 0.426 |
| Notch_signaling_pathway_[hsa04330] | 64 | 0.344 | 196 | 0.310 |
| Oxytocin_signaling_pathway_[hsa04921] | 154 | 0.376 | 456 | 0.373 |
| p53_signaling_pathway_[hsa04115] | 710 | 0.301 | 9841 | 0.217 |
| Phosphatidylinositol_signaling_system_[hsa04070] | 290 | **0.652** | 1892 | **0.625** |
| Phospholipase_D_signaling_pathway_[hsa04072] | 106 | 0.388 | 264 | 0.386 |
| PI3K-Akt_signaling_pathway_[hsa04151] | 308 | 0.403 | 1956 | 0.364 |
| PPAR_signaling_pathway_[hsa03320] | 968 | 0.643 | 4608 | 0.159 |
| Prolactin_signaling_pathway_[hsa04917] | 170 | 0.380 | 884 | 0.345 |
| Rap1_signaling_pathway_[hsa04015] | 570 | 0.485 | 5707 | 0.397 |
| Ras_signaling_pathway_[hsa04014] | 479 | 0.413 | 3337 | 0.412 |
| Relaxin_signaling_pathway_[hsa04926] | 170 | 0.395 | 558 | 0.392 |
| RIG-I-like_receptor_signaling_pathway_[hsa04622] | 101 | 0.442 | 355 | 0.427 |
| Sphingolipid_signaling_pathway_[hsa04071] | 115 | 0.405 | 301 | 0.400 |
| T_cell_receptor_signaling_pathway_[hsa04660] | 176 | 0.487 | 676 | 0.478 |
| TGF-beta_signaling_pathway_[hsa04350] | 75 | 0.414 | 170 | 0.392 |

| | | | | |
|---|---|---|---|---|
| Thyroid_hormone_signaling_pathway_[hsa04919] | 207 | 0.338 | 848 | 0.344 |
| TNF_signaling_pathway_[hsa04668] | 84 | 0.350 | 172 | 0.343 |
| Toll_and_Imd_signaling_pathway_[dme04624] | 146 | 0.460 | 409 | 0.457 |
| Toll-like_receptor_signaling_pathway_[hsa04620] | 343 | 0.408 | 1346 | 0.391 |
| VEGF_signaling_pathway_[hsa04370] | 84 | 0.376 | 278 | 0.358 |
| Wnt_signaling_pathway_[hsa04310] | 426 | 0.448 | 2529 | 0.435 |

Supplementary Table S17. The results of $N_3$, $N_4$, $H_{3R}$ and $H_{4R}$ for cellular processes.

| Cellular processes | $N_3$ | $H_{3R}$ | $N_4$ | $H_{4R}$ |
|---|---|---|---|---|
| Adherens_junction_[hsa04520] | 362 | 0.539 | 2273 | 0.524 |
| Apoptosis_[hsa04210] | 567 | 0.463 | 3214 | 0.450 |
| Cell_cycle_[hsa04110] | 205 | 0.419 | 780 | 0.402 |
| Cellular_senescence_[hsa04218] | 186 | 0.408 | 595 | 0.369 |
| Focal_adhesion_[hsa04510] | 359 | 0.474 | 1759 | 0.455 |
| Gap_junction_[hsa04540] | 97 | **0.361** [*] | 303 | **0.336** [*] |
| Necroptosis_[hsa04217] | 336 | 0.556 | 2471 | 0.525 |
| Regulation_of_actin_cytoskeleton_[hsa04810] | 193 | 0.449 | 727 | 0.409 |
| Signaling_pathways_regulating_pluripotency_of_stem_cells_[hsa04550] | 85 | **0.614** [𝔍] | 182 | **0.532** [𝔍] |

The symbols [*] and [𝔍] denote the minimum and maximum respectively.

Supplementary Tables S18. The ranges of $H_{3R}$ and $H_{4R}$ for cancer networks, STN and cellular processes.

| Network type | range of $H_{3R}$ | range of $H_{4R}$ |
|---|---|---|
| cancer networks | 0.323 ~ 0.493 = 0.170 | 0.273 ~ 0.483 = 0.210 |
| STN | 0.120 ~ 0.652 = 0.532 | 0.034 ~ 0.625 = 0.622 |
| cellular processes | 0.361 ~ 0.614 = 0.253 | 0.336 ~ 0.532 = 0.196 |
